# Characterizing the role of mitochondrial dynamics during *Drosophila* convergent extension using NADH fluorescence lifetime imaging

**DOI:** 10.64898/2025.12.30.693754

**Authors:** Maria Espana-Pena, Alan Woessner, Colten Nichols, Kyle P. Quinn, Adam C. Paré

## Abstract

Mitochondria are dynamic organelles that can fragment or fuse to support different bioenergetic demands (e.g., glycolysis vs. oxidative phosphorylation) and distinct cell behaviors (e.g., mitosis or migration). While the role of mitochondrial dynamics in wound healing and metabolic disorders has received significant attention, the role of mitochondrial fission and fusion during normal embryonic development is less well understood––in part due to the difficulty of studying such processes *in vivo*. Combined with the depth-resolved imaging capabilities of multiphoton microscopy, fluorescence lifetime imaging (FLIM) of the mitochondrial cofactor NADH can be used to simultaneously visualize mitochondrial network morphology and infer certain aspects of cellular bioenergetics (e.g., glycolysis vs. oxidative phosphorylation) in a label-free, non-invasive manner. Here we demonstrate that NADH FLIM can be used to accurately track the subcellular localization and topology of mitochondrial networks in live *Drosophila* embryos. We used this technique to assess whether cells show changes in NADH lifetime during convergent extension (CE)––a conserved process of tissue remodeling in which thousands of germband cells undergo coordinated intercalation to drive elongation of the head-to-tail axis. Contrary to our expectations, we did not observe significant changes in NADH lifetime or network appearance during CE in wild-type embryos, suggesting that germband cells do not need to alter their baseline metabolism to fuel cell intercalation during normal development. To directly assess the role of mitochondrial fission and fusion during CE, we used RNA interference to disrupt the fission mediator *Drp1* and the fusion mediator *Opa1*. Consistent with expectations, inhibiting mitochondrial fission in *Drp1-*knockdown embryos led to hyper-fused networks and significantly longer NADH lifetimes, indicating a shift towards oxidative phosphorylation. Conversely, inhibiting mitochondrial fusion in *Opa1-*knockdown embryos led to more hyper-fragmented networks and significantly shorter NADH lifetimes, indicating a shift towards glycolysis. Interestingly, inhibiting either fission or fusion altered tissue elongation and greatly increased the rate of cell intercalation errors, suggesting that a precise network topology is required for proper CE. We hypothesize that the CE defects in *Drp1*-knockdown embryos are primarily due to incorrect basal subcellular localization of mitochondria, whereas the CE defects in *Opa1*-knockdown embryos are due to deficient ATP and/or ROS production. These experiments demonstrate the utility of FLIM-based applications for characterizing the role of mitochondria during normal embryonic development, which could yield a better understanding of the metabolic underpinnings of various pathologies that involve epithelial remodeling, including spina bifida, defective wound healing, and cancer metastasis.

## INTRODUCTION

Mitochondrial networks are highly complex organelles that can fuse, fragment, or move to different regions of the cell to support a wide variety of cellular processes. Beyond their central importance to ATP generation and metabolism, it is now clear that mitochondria also play critical roles as local signaling centers and sources of small molecules to effect changes in cell fate, behavior, and morphology (Madan et al., 2022; Onishi et al., 2021; Palikaras & Tavernarakis, 2014; Westermann, 2012). The particular architecture of a cell’s mitochondrial network at any point in time is a balance between fission and fusion, which is largely controlled by the activity of dynamin-related protein 1 (Drp1) and optic nerve atrophy 1 (Opa1), respectively (Chen et al., 2023; Liesa & Shirihai, 2013; Mishra & Chan, 2016; Ponte et al., 2020). Studies in multiple organisms have demonstrated that disrupting mitochondrial dynamics––the complex interplay between fission and fusion that shapes mitochondrial network morphology (Kiefel et al., 2006; Kyriakoudi et al., 2021; Mishra & Chan, 2016; Youle & Van Der Bliek, 2012; Zhao et al., 2013)––leads to defects in wound healing (Dhall et al., 2014; Duan et al., 2020; Jones & Quinn, 2021; Lowell & Shulman, 2005; Mehrvar et al., 2019; Miskolci et al., 2022; Moriyama et al., 2025; Sood et al., 2015; Taner et al., 2024; Xu et al., 2022; Xu & Chisholm, 2014), indicating that mitochondrial processes can directly affect cell morphology and tissue remodeling. However, the importance of mitochondrial localization and dynamics during normal embryonic development, particularly in the context of tissue remodeling, has received relatively little study.

A popular model system for studying tissue remodeling is the fruit fly *Drosophila melanogaster*, as embryos are easy to obtain, they are amenable to genetic manipulation, and the cell rearrangements that drive the early events of embryogenesis occur on the surface of the embryo, making them easy to visualize by microscopy. Extension of the head-to-tail axis offers an interesting opportunity to study how a tissue––in this case the germband neuroectoderm––transitions from a static state to one in which hundreds of cells undergo coordinated and rapid cell intercalation to drive tissue elongation. This process of cell intercalation causing tissue elongation is known as convergent extension (CE), and it is a highly conserved general mechanism for shaping tissues and organs in all animals (Paré et al., 2014; Shindo, 2018; Walck-Shannon & Hardin, 2013). It is possible to capture the entire process of CE in living *Drosophila* embryos by microscopy because it occurs on a very short timescale compared with vertebrates, with the majority of cell intercalation occurring over the course of 30 minutes (Irvine & Wieschaus, 1994; Kasza et al., 2014; Paré et al., 2019; Tamada & Zallen, 2015). Studies using conventional fluorescent confocal imaging have revealed much about the biomechanical processes that drive cell intercalation, which are caused by changes in the subcellular localization and activity of actin, myosin, and adherens junction proteins to shrink existing apical cell-cell contacts or build new ones (Lecuit, 2004; Zallen & Wieschaus, 2004)––energy-intensive processes that presumably require increased ATP levels, although this has never been demonstrated. Most of these proteins are also directly involved in cell movements during wound healing (Scepanovic et al., 2021), which are also linked to mitochondrial dynamics. Finally, it has been shown that mitochondria relocate from the basal to the apical cell surface immediately before CE in the *Drosophila* neuroectoderm (Chowdhary et al., 2017, 2020). Therefore, we hypothesized that there might be distinct changes in mitochondrial localization, morphology and/or metabolism as cells begin to reorganize within this tissue during CE.

Label-free multiphoton microscopy has become a useful approach to visualize metabolic changes and capture cellular and tissue-level morphological and biochemical features in 3D without the addition of exogenous stains or dyes (Georgakoudi & Quinn, 2023). Specifically, two-photon excited fluorescence (TPEF) microscopy can detect the natural fluorescence of the reduced form of nicotinamide adenine dinucleotide (NADH), a key metabolic cofactor that transfers electrons between the tricarboxylic acid (TCA) cycle with the electron transport chain (ETC). NADH shifts between a free or protein-bound state when transferring electrons, which affects the time that NADH spends in an excited state prior to a return to its ground level with the emission of a photon. When TPEF is combined with fluorescence lifetime imaging (FLIM), we can use these time-resolved measurements to determine the proportion of free-to-bound. A greater proportion of free NADH has been associated with more glycolysis and carbon substrate catabolism with short lifetimes (0.3-0.4 ns), while a greater proportion of bound NADH has been associated with a moderate increase in oxidative phosphorylation and relatively long lifetimes (1.9-5.7 ns) (Georgakoudi & Quinn, 2023; Kolenc & Quinn, 2019). The fluorescence lifetime of NADH has been widely used in studies of stem cell differentiation, wound healing and its treatment, neuroscience, aging, infectious and non-infectious disease (Ameer-Beg et al., 2020; Jones & Quinn, 2021; Kolenc & Quinn, 2019; Morrow et al., 2024; Rico-Jimenez et al., 2020; Snyder et al., 2023; Suhling et al., 2015; Torrado et al., 2024).

However, there have been relatively few studies where NADH FLIM has been applied to developmental biology of healthy tissues (Ma et al., 2019; Sanchez et al., 2018; Seidler et al., 2020; Supatto et al., 2005, 2009, 2011).

This study investigated whether TPEF and NADH fluorescence lifetime imaging can be used to non-invasively assess the bioenergetics of tissue remodeling during convergent extension in the *Drosophila* embryo in vivo. To address this question, we performed high-speed multiphoton FLIM before, during and after convergent extension (∼45 minutes) in live embryos. NADH FLIM measurements were sensitive to alterations in mitochondrial function and localization following inhibition of mitochondrial fission and fusion. To our knowledge, this is the first in vivo assessment of NADH lifetime dynamics during convergent extension in wild-type, fission-impaired, and fusion-impaired *Drosophila* embryos. The successful implementation of this approach to characterize convergent extension in vivo highlights the potential of NADH FLIM as a powerful tool for probing tissue remodeling, with applications that may extend to additional developmental processes in *Drosophila* and other model organisms.

## METHODS

### *Drosophila* husbandry, stocks, and crosses

Adult F1 flies used for experiments were reared on a standard food mixture containing molasses, cornmeal, yeast, malt, agar, and preservatives (Tegosept and propionic acid) and maintained at 25°C; parental stock lines were maintained between 18–25°C. Embryonic stages used in this study were as follows (at 25°C): stage 5 ≈ 2.0–2.5 h AEL, stage 6 ≈ 2.5–3.0 h AEL, stage 7 ≈ 3.0–3.25 h AEL, and stage 8 ≈ 3.25–4.0 h AEL. The sex of the embryos was not considered relevant and was not determined in this study.

The following fly stocks were used: 1) *yellow white* wild-type control (*y[1]*, *w[1]*) (BDSC #1495); 2) *αtubulin–Gal4[13]; αtubulin–Gal4[4]* maternal driver (referred to as *mat13;4*) (gift of Drs. Richa Rikhy and Girish Ratnaparkhi); 3) *αtubulin–Gal4[67]; αtubulin–Gal4[15]* maternal driver (referred to as *mat67;15*) (gift of Dr. Jennifer Zallen); 4) UAS–Drp1 RNAi (*y[1], v[1]; P{y[+t7.7] v[+t1.8]=TRiP.HMC03230}attP40*) (BDSC #51483); 5) UAS–Opa1 RNAi (*w[*]; Bl[1]/CyO; P{w[+mC]=UAS-Opa1.RNAi.CDS}3*) (BDSC #67159); 6) Mito–GFP mitochondrial marker (*w[*]; P{w[+mC]=sqh-EYFP-Mito}3*) (BDSC #7194); 7) gap43–mCherry cell membrane marker (gift of Dr. Adam Martin).

For *in vivo* confocal imaging of mitochondria and convergent extension, we created a stock containing αtubulin–Gal4[67] on chromosome II, plus gap43–mCherry and Mito–GFP recombined on chromosome III (referred to as *mat67; gap43, Mito*). To create knockdown embryos, we crossed *mat67; gap43, Mito* virgin homozygous females to UAS–Drp1 RNAi or UAS–Opa1 RNAi homozygous males to create transheterozygous F1 offspring; for controls, *yellow white* males were used instead of UAS RNAi males. F1 adults were placed in a cage, and F2 embryos were collected and imaged (below). For RNAi knockdown and fluorescent transgenes in the early embryo, maternal expression was considered most important, and the exact zygotic genotypes of the F2 embryos were not determined.

For live multiphoton FLIM microscopy and fixed confocal microscopy, no fluorescent transgenes were present. To create *Drp1* knockdown embryos, we crossed *mat67;15* (for stronger knockdown) or *mat13;4* (for weaker knockdown) virgin homozygous females to UAS–Drp1 RNAi homozygous males. *Opa1* knockdown embryos were created by crossing *mat67;15* virgin homozygous females to UAS–Opa1 RNAi homozygous males. For the controls, *yellow white* males were used instead of UAS RNAi males.

### Immunostaining of fixed embryos

For embryo collections, F1 adults were placed in ventilated cages with apple-juice agar plates and allowed to lay for ∼7 h at 25°C. Embryos were dechorionated in 1:1 bleach:water for 3 min and rinsed well. Dechorionated embryos were fixed in a 1:1 mixture of heptane:18% formaldehyde (VWR, Cat. #76180-690) in 1x phosphate buffered saline (PBS) and shook for 20 min at room temperature. Fixed embryos of the desired stages were manually devitellinized with a fine needle (Parveen et al., 2023). Immunohistochemistry was performed using standard procedures. In brief, embryos were washed in PBS+0.1% Triton X-100 for 20 min, blocked with 10% BSA in PBS+0.1% Triton X-100 for 1 h, incubated with primary antibodies overnight, washed with PBS+0.1% Triton X-100 for 2 h, incubated with secondary antibodies for 1 h, washed with PBS+0.1% Tween20 for 2 h, and mounted in ProLong Gold antifade mountant with DAPI (Thermo Fisher #P36935). Primary antibodies used were: mouse anti-ATP5A1 (1:300, Thermo Fisher #439800), guinea pig anti-Par-3/Bazooka (1:300, gift of Dr. Jennifer Zallen), and rabbit anti-β-catenin/Armadillo (1:300, gift of Dr. Jennifer Zallen). Secondary antibodies used were: goat anti-guinea pig Alexa647 (1:500, Thermo Fisher #A-21450), goat anti-rabbit Alexa546 (1:500, Thermo Fisher #A-11035), and goat anti-mouse Alexa488 (1:500, Thermo Fisher #A11029).

### *In vivo* confocal imaging of convergent extension and mitochondrial networks

F2 embryos were collected and dechorionated as described above. Embryos were placed in halocarbon oil 700 (HO700) (Millipore Sigma #H8773) using a fine paintbrush. Blastoderm (stage five) embryos were identified using a dissection microscope and mounted in HO700 between an oxygen-permeable membrane (YSI Life Sciences #66155) and a #1.5 glass coverslip in a custom mounting slide. Embryos were imaged on an Axio Observer 7, Zeiss LSM900 confocal microscope with a Plan-Apochromat 40x/1.3 NA oil-immersion objective (Zeiss). Non-saturated 8-bit image stacks were collected (Zen 3.0 software) from the ventrolateral apical embryo surface at 1x zoom (1024 x 1024 pixels) with a 1 AU pinhole and a 0.45–0.5 µm interval between slices. Maximum intensity projections were created using ImageJ/Fiji. To track cell movements and mitochondria, non-saturated 8-bit image stacks of gap43–mCherry or Mito–GFP signal were collected from the ventrolateral apical embryo surface every 30 s at 0.8-1x zoom (768 x 600 pixels, 0.25 µm/pixel), with 2x averaging, a 1 AU pinhole and a 0.45–0.5 µm interval between slices. Using Fiji/ImageJ, three apical slices from each timepoint were projected (average intensity) to produce the final movies. To quantify tissue elongation, we identified groups of four cells in a rectangular arrangement that could be tracked over the course of the entire movie, and we calculated the change in aspect ratio (width/height) every 5 min. To account for slight differences in shape between the cell groups, all aspect ratio measurements were normalized to the original aspect ratio at t=0. To register the different movies in time, t=0 was set as 10 frames before the first cell entirely flowed off the posterior edge of the movie frame, which closely correlates with the onset of stage 7 (closure of the ventral midline and the beginning of cell intercalation) in wild-type embryos. To determine edge formation errors, ∼50 interfaces were visually examined per movie (n=3 per genotype) for a total of ∼150 interfaces per genotype. The fate of each edge was categorically scored as resolving either correctly or incorrectly. Correctly resolved interfaces were classified as “productive”, whereas incorrectly resolved interfaces were further categorized as “no edge formation” (edge remained unresolved), “wrong direction” (vertical resolution instead of horizontal), or “no shrink” (no change in edge length over time).

### *In vivo* two-photon-excited fluorescence microscopy and FLIM

F2 embryos were collected, dechorionated, and mounted as described above. Mounted embryos were placed in a temperature-controlled microincubator (DH-35IL; Warner Instruments) and maintained at 25 °C during imaging. Acquisition was performed using a Bruker Investigator+ multi-photon microscope (Bruker) coupled with an ultrafast Ti:Sapphire pulsed laser (InsightX3; Spectra-Physics) to acquire TPEF (755 nm ex./460 nm em.). Single plane and Z-stack images were acquired at neuroectoderm regions, using a 25x/1.10 NA Apochromatic LWD water-immersion objective (Nikon). Time-correlated single-photon counting was performed using a Time Tagger Ultra (Swabian). Images were acquired with an optical zoom of 7-8x (256x256 pixels, 0.29 µm/pixel) covering a tissue region of approximately 65 µm x 65 µm, encompassing ∼150 cells. The laser repetition rate was 12.5 ns, with an integration time of 10 seconds per image, and the incident laser power maintained at or below 5 mW at all depths to ensure embryo survival. To offset lower photon counts due to the shorter integration time and lower laser power, we increased the pixel dwell time to 3.6 µs and applied additional binning post-processing to increase our per-pixel photon count to >8,000 total photons. FLIM single plane images were acquired first, followed by Z-stacks from the top of the epithelium (most apical) to a depth of ∼20 μm (most basal) in increments of 0.5 μm, for each field of view.

### FLIM data processing

Time-correlated single photon counting was used to record the timing of individual photons detected relative to the laser pulses. These photon arrival times were used to create time decay histograms of NADH fluorescence at each pixel. The raw decay data were imported into SPCImage 8.3 software (Becker & Hickl GmbH, Germany) to: 1) deconvolve the measured decay and the instrument response function of the system that was acquired using the second harmonic signal of a BBO crystal, and 2) fit the lifetime decay data to a bi-exponential model,

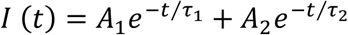

where 𝜏_1_ represents the lifetime of free NADH, 𝜏_2_ represents the lifetime of bound NADH, and 𝐴_1_ and 𝐴_2_represent the relative proportion of free and bound NADH, respectively (Kolenc & Quinn, 2019). A weighted least squares model of incomplete multiexponential decay was used. To increase the signal-to-noise ratio, pixel binning was used to ensure at least 8,000 total photons, with a maximum bin size of 10 (21x21 pixels). Color maps were acquired from SPCImage and standardized (0.1 ns = red, 3.0 ns = blue) across all images to show mean lifetime (τ_m_) per pixel.

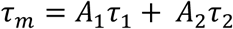

After fitting was completed and color ranges were set, the data matrices representing fluorescence lifetime components (A1, A2, 𝜏_1_, and 𝜏_2_) were exported out of SPCImage and mean NADH lifetimes, as well as free-to-bound NADH contributions were calculated using a custom-made MATLAB script.

## Statistical analysis

Two-way ANOVA (or mixed model) and Tukey’s multiple comparison tests were used to assess differences in NADH fluorescence lifetime across developmental stages (5–8) and between control embryos (n=6), Drp1 knockdowns (n=3) and Opa1 knockdowns (n=4). To evaluate edge formation errors, confocal movies (n=3/genotype) were independently analyzed by two investigators. A total of ∼150 vertices/genotype were categorically scored as previously described. The proportion of identified errors was then calculated as a percentage and averaged across the two investigators. Then, Pearson’s chi square tests were used to determine significant changes between observed and expected data for edge formation errors among genotypes. All analyses were performed using GraphPad Prism 10.6.1.

## RESULTS

### NADH autofluorescence reveals the subcellular morphology and localization of mitochondria in unlabeled living *Drosophila* embryos

To assess the practicality of using NADH autofluorescence for visualizing mitochondria in unlabeled living embryos using TPEF microscopy, we chose to analyze the germband neuroectoderm in the early *Drosophila* embryo. Cells in this tissue are initially arranged in a static hexagonal grid but then begin to rapidly intercalate at the onset of gastrulation to drive tissue elongation via convergent extension (CE) (Irvine & Wieschaus, 1994). Cell intercalation is driven by dramatic changes in the subcellular localization and activity of actin and myosin to contract specific apical cell-cell contacts (Lecuit, 2004; Zallen & Wieschaus, 2004), which presumably requires increased ATP levels to fuel these processes. Consistent with this, after cellularization of the blastoderm is complete (stage 5), mitochondria relocate from the basal to the apical cell surface immediately before intercalation begins (Chowdhary et al., 2017). Therefore, we hypothesized that there might be distinct changes in mitochondrial architecture and/or metabolic profiles as cells begin to reorganize within the tissue.

Stage-5 blastoderm embryos (2.0–2.5 h AEL) were manually selected and mounted between a coverslip and a gas-permeable membrane in halocarbon oil and maintained at 25°C throughout stages 5–8 for ∼30 min of imaging (Figure 1A). NADH lifetime data was collected from single x-y planes or z-stacks (n=4 FOV per embryo) from the ventrolateral neuroectoderm, with z-stacks spanning from the most apical surface to a depth of approximately 22 µm (Figure 1C). Acquisition parameters (dwell-time, integration time, and laser power) were optimized for FLIM, and images of NADH subcellular localization were generated for stages 5–8 (see Methods). As a comparison, we performed immunofluorescence with a standard mitochondrial marker (ATP5A1) in fixed embryos of comparable stages (Figure 1B). We found that NADH intensity (grayscale) and conventional mitochondrial immunofluorescence signals showed very similar spatial localization patterns before (stages 5 & 6), during (stage 7) and after CE (stage 8) in control embryos (n=6). During stage 5, mitochondria were predominantly perinuclear, consistent with previous reports describing fragmented mitochondria moving basally to apically during cellularization (Chowdhary et al., 2020; Madan et al., 2022). During stage 6, mitochondria were predominantly located in the apical region of the cell (0–7 µm), but were often excluded from the most medial region, directly above the nucleus, giving the networks a slight donut-like appearance (Figure 1C). During stage 7, which encompasses the fast phase of CE when cells are actively intercalating (∼25 min), the networks lose their donut-like appearance and become more homogeneously localized within the apical region. During stage 8, mitochondrial networks begin to adopt different morphologies in different cells, presumably linked to asynchronous cell divisions that begin to occur in this tissue after intercalation (Edgar & O’Farrell, 1989). Consequently, we confirmed that *in vivo* imaging of mitochondrial network architecture in unlabeled *Drosophila* embryos using TPEF is non-invasive, as it does not interfere with normal mitochondrial localization, and can be used to resolve subcellular details comparable with conventional confocal immunofluorescence in fixed tissues.

**Figure 1.**
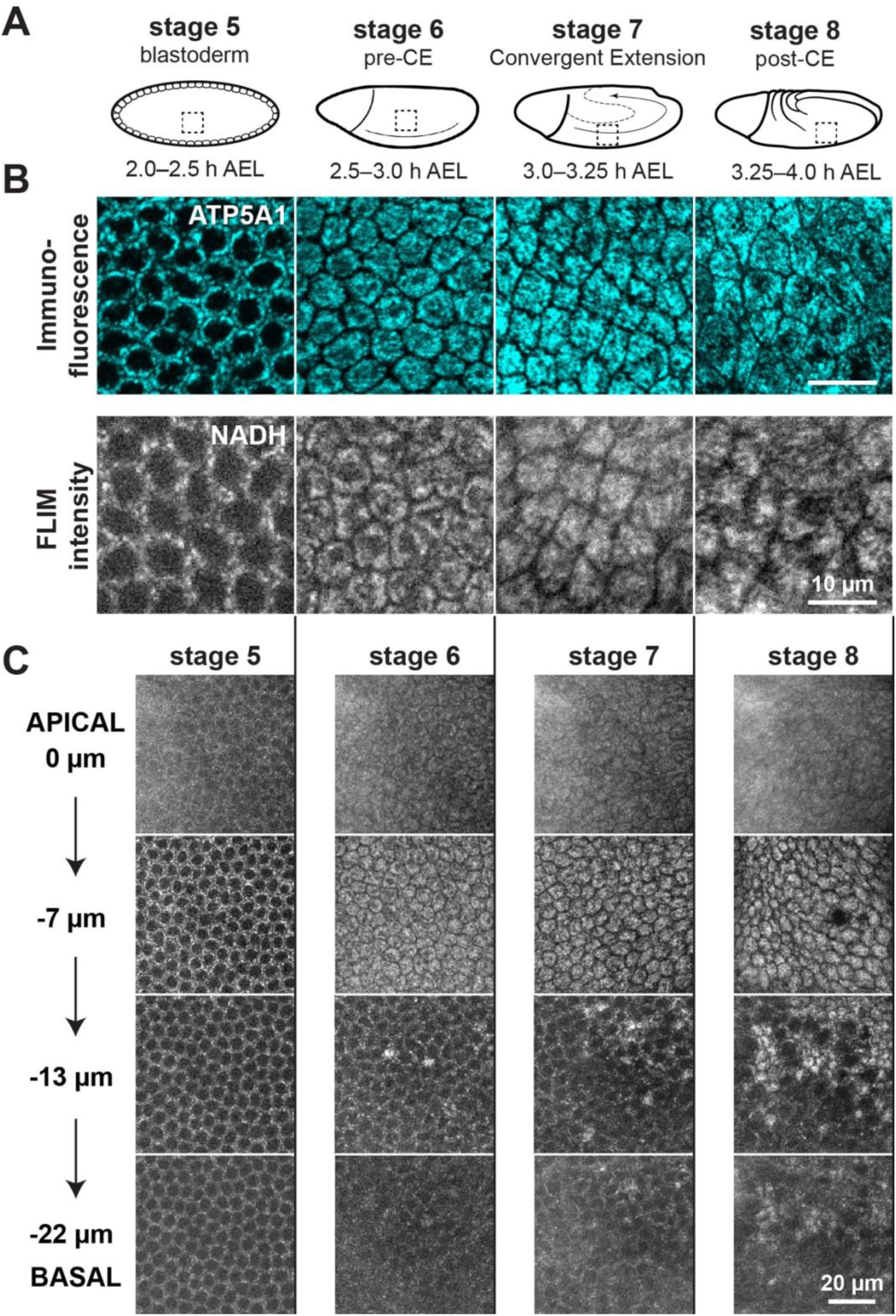
Visualization of subcellular mitochondrial morphology and localization using NADH autofluorescence during *Drosophil*a convergent extension. A) Schematics showing stages of *Drosophila* embryogenesis around convergent extension: stage 5 – monolayer epithelium cellularizes to form the blastoderm; stage 6 – cellularization is complete and gastrulation begins, but germband cells are static; stage 7 – active cell intercalation and convergent extension of the germband; stage 8 – intercalation largely complete, cell divisions begin. AEL: After Egg Laying. **B)** Top panels: fixed wild-type embryos stained with a mitochondrial marker (ATP5A1, cyan) and imaged using standard laser-scanning confocal microscopy. Bottom panels: live, unlabeled wild-type embryos, imaged using two-photon excitation fluorescence microscopy (340–350 nm ex./460–470 nm em.) to visualize NADH (grayscale). **C)** Mitochondrial localization along the apical-basal cell axis in wild-type embryos. Slight glare from extraembryonic vitelline membrane was used as apical reference point (0 µm).

### Genetic perturbation of mitochondrial dynamics alters network localization and organization in the *Drosophila* neuroectoderm

Next, we sought to determine the role of mitochondrial dynamics during convergent extension. In particular, we were interested in determining whether there are changes in network architecture over the course of CE, and whether these changes are necessary for CE to occur. The particular architecture of a cell’s mitochondrial network at any point in time is a balance between fission and fusion, which is largely controlled by the activity of dynamin-related protein 1 (Drp1) and optic nerve atrophy 1 (Opa1), respectively (Chen et al., 2023; Liesa & Shirihai, 2013; Mishra & Chan, 2016; Ponte et al., 2020). It is not possible to completely remove the function of these proteins, as this would be lethal to the embryo. Therefore, we disrupted fission and fusion in the early embryo by using RNA interference to knock down the expression levels of *Drp1* and *Opa1*. Nearly all proteins and RNAs in the very early embryo are maternal in origin, meaning they were produced by the mother’s germline cells and transported into the egg. Therefore, we used the UAS-Gal4 system to drive maternal expression of double-stranded RNAs complementary to *Drp1* and *Opa1* messenger RNAs to target them for destruction in the egg.

To confirm the efficacy of our knockdowns, we used confocal microscopy and immunofluorescence to visualize mitochondrial network architecture (ATP5A1) and cell morphology using an apical adherens junction marker (β-catenin/Armadillo) in stage 5–8 embryos (Figure 2). In cells from control embryos, mitochondrial networks moved from the perinuclear region (stage 5) to the apical region (stages 6–8), as described above, but individual mitochondria could not be clearly resolved, making it difficult to determine whether the network as a whole is in a fused or fragmented state (Figure 2A). Inhibition of the fission mediator Drp1 had a number of effects on mitochondrial localization and morphology at all stages analyzed. In *Drp1*-knockdown embryos, we observed very little mitochondrial signal in perinuclear regions during stage 5 or in the apical region during stage 6, and the few signals that were present were small and punctate (Figure 2B). By stage 7, the majority of mitochondria did reach the apical region, but they showed a distinct morphology compared with controls, often appearing more thread-like, consistent with hyper-fused networks (Mitra et al., 2012; Smirnova et al., 2001) (Figure 2B). During stage 8, these differences were even more apparent, with mitochondria forming large clumps localized to one side of the nucleus (Figure 2B). By contrast, inhibition of the fusion mediator Opa1 had little effect on mitochondrial localization but had a distinct effect on mitochondrial morphology. In *Opa1*-knockdown embryos, mitochondria showed the proper perinuclear to apical migration, but the signals were much more punctate than in controls, consistent with hyper-fragmented networks (Figure 2C). Therefore, as inhibition of fission and fusion both led to networks with distinct morphologies compared with controls, we conclude that during wild-type CE, mitochondrial networks exist in an intermediate state between fully fused and fully fragmented. Furthermore, we conclude that there are not dramatic changes in mitochondrial network morphology before, during, or after CE.

**FIGURE 2.**
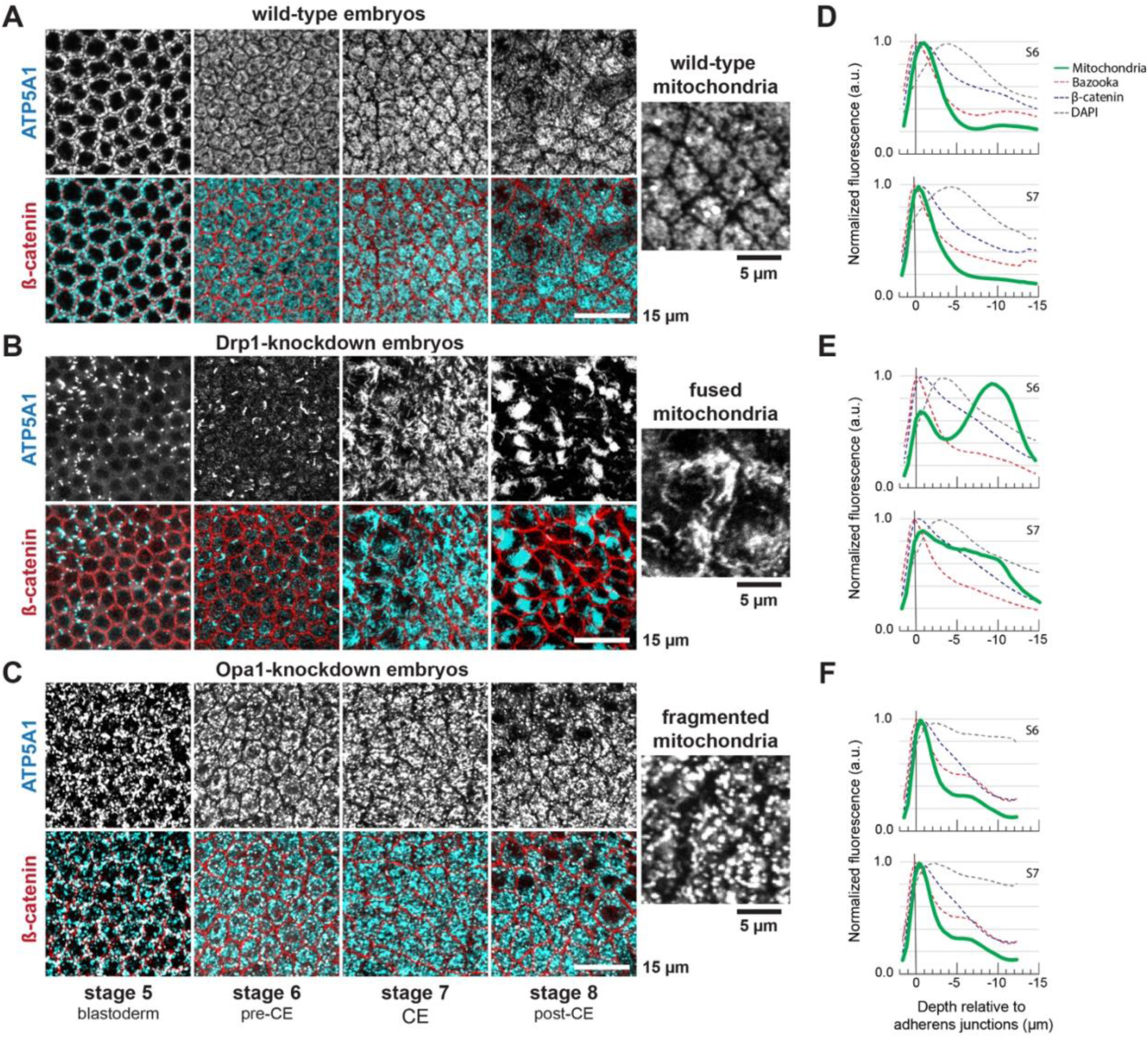
Inhibiting mitochondrial fission and fusion alters mitochondrial network architecture and localization during convergent extension. Visualization of mitochondria and cell morphology using immunofluorescence labeling of complex IV (ATP5A1, grayscale in top panels, cyan in bottom panels) and adherens junctions as a membrane marker (β-catenin, red in bottom panels) in control embryos **(A)**, Drp1-knockdown embryos **(B)**, and Opa1 knockdown embryos **(C)**. Right panels show zoomed in views of mitochondria from stage-7 embryos. Knockdown was achieved through genetically encoded RNA interference using maternally provided double-stranded RNAs targeting the *Drp1* and *Opa1* messenger RNAs. Fluorescence intensity across apical, perinuclear and basal regions before (S6 = stage 6) and during (S7 = stage 7) convergent extension, using labels ATP5A1 for mitochondria (green solid line), β-catenin/Armadillo (blue dotted line) and Bazooka/Par-3 (red dotted line) for adherens junctions, and DAPI for nuclei (gray dotted line). The intensity peak of the marker for apical adherens junction protein Bazooka was used as a reference point (0 µm) in control embryos **(D)**, Drp1-knockdown embryos **(E)**, and Opa1-knockdown embryos **(F)**.

To better characterize the differences in mitochondrial apical-basal localization between control and *Drp1-*knockdown embryos, we collected fluorescence data spanning the apical, perinuclear, and basal regions prior to CE (stage 6) and during CE (stage 7). Mitochondria were marked with ATP5A1, apical adherens junctions were marked with Par-3/Bazooka and β-catenin/Armadillo, and nuclei were marked with the fluorescent DNA stain DAPI (Figure 2D-F). Using the peak of Par-3 fluorescence intensity as a reference point (0 µm), in control embryos, the majority of mitochondria were located just below the adherens junctions at stage 6 (-0.90 µm) and at stage 7 (-0.45 µm), with very little mitochondrial signal extending into the perinuclear or basal regions of the cell (Figure 2D). *Opa1-*knockdown embryos showed very similar mitochondrial distributions to controls, with signals peaking just below the adherens junctions during stage 6 (-0.45 µm) and stage 7 (-0.45 µm) (Figure 2F). By contrast, *Drp1-*knockdown embryos showed a bimodal distribution of mitochondria during stage 6, with a small peak just below the adherens junctions (-0.45 µm) and a larger peak localized below the nucleus (-9.0 µm). During stage 7, the highest peak of mitochondrial signal is now located just below the adherens junctions (-0.45 µm), but a significant portion of the signal can still be seen in the perinuclear and basal regions of the cell (Figure 2E). Therefore, we conclude that inhibiting mitochondrial fission disrupts the proper subcellular localization of mitochondria, causing networks to remain in the basal region of the cell. This is consistent with defective migration of hyper-fused mitochondria from the basal to the apical cell surface observed in *Drp1*-knockdown embryos during stage 5 (Chowdhary et al., 2017), and we show here that these defects persist at least into stage 7.

### Inhibition of mitochondrial fission and fusion cause defects in cell intercalation during CE

We next asked whether disrupting mitochondrial dynamics interferes with convergent extension. To assess the overall progression of CE, we collected time-lapse images of germband cells undergoing intercalation (stage 5 through stage 8), and then quantified tissue elongation by calculating the change in aspect ratio (Δ aspect ratio) of a rectangular group of cells on the lateral embryo surface over time. In control embryos, the aspect ratio (length/height) approximately doubled (Δ aspect ratio = 1.97) after 15 minutes of cell intercalation, and these changes were reproducible between embryos (std. dev. = 0.15), consistent with previous studies (Paré et al., 2019; Zallen & Wieschaus, 2004). We also confirmed that the surface area of the group of cells stayed essentially constant during those 15 minutes, demonstrating that CE is driven by cell rearrangements, not tissue growth.

When knocking down *Drp1* and *Opa1* using RNAi, we observed a large range of phenotypes with respect to gross cell morphology immediately prior to CE (stage 6): some knockdown embryos appeared indistinguishable from controls, whereas other knockdown embryos displayed highly atypical cell morphologies (e.g., very large cells or cells with multiple nuclei). We hypothesized that these different phenotypes were due to variations in the efficacy of RNAi from embryo to embryo, with very strong knockdown of *Drp1* and *Opa1* causing disruptions during very early stages of development that impaired cellularization of the epithelium (stage 5). Therefore, we only analyzed embryos that appeared wild type at the start of imaging (stage 6), which were likely embryos with mild gene knockdown. In *Drp1-*knockdown (Δ aspect ratio = 1.82) and *Opa1-*knockdown embryos (Δ aspect ratio = 2.03), the average change in aspect ratio approximately doubled after 15 minutes of intercalation, similar to controls (not significant by t-test or ANOVA). However, unlike controls, the variation in tissue elongation from embryo to embryo was quite large (*Drp1*-knockdown std. dev = 0.64, *Opa1-*knockdown std. dev. = 1.32), with some embryos elongating less than on controls, and others elongating more than controls. These results suggest that inhibiting both mitochondrial fission and fusion can have effects on tissue elongation during CE, although the associated effects in tissue geometry are highly variable.

Considering that tissue elongation is a direct result of cell intercalation (Irvine & Wieschaus, 1994) we next decided to analyze the cell rearrangements that drive CE. The most common mode of cell intercalation (T1 or neighbor exchange) involves a group of four cells that initially share a vertical cell-cell interface (Figure 3A). Mediated by planar polarized actomyosin and adherens junctions, vertical interfaces contract and disappear, temporarily bringing all four cells into contact, and a new interface is then formed in the horizontal direction, causing cells to intercalate and contributing to productive tissue elongation. In control embryos, 80% of vertical interfaces underwent productive transitions to horizontal interfaces, whereas in 20% of cases we observed one of three error types: 1) the initial vertical interface did not contract (4%), 2) the 4-cell vertex never formed a new interface (8%), or 3) the new interface formed in the vertical (rather than horizontal) orientation (8%, Figure 3B). We observed many more intercalation errors compared with controls in both *Drp1-*knockdown embryos (61% vs. 20%, p < 0.0001) and *Opa1-*knockdown embryos (74% vs. 20%, p < 0.0001), with most vertical interfaces failing to undergo productive transitions in both backgrounds (Figure 3B). For the *Drp1*-knockdown embryos, the errors we observed were primarily type-2 (no edge formation) and type-3 (unproductive orientation) errors. In *Opa1*-knockdown embryos, all three error types were observed in roughly equal proportions, although the most notable increase relative to controls was in type-1 errors (no interface contraction). These data demonstrate that inhibiting mitochondrial fission or fusion can strongly disrupt cell intercalation and that distinct steps of intercalation may be differentially affected by *Drp1* or *Opa1* knockdown.

**Figure 3.**
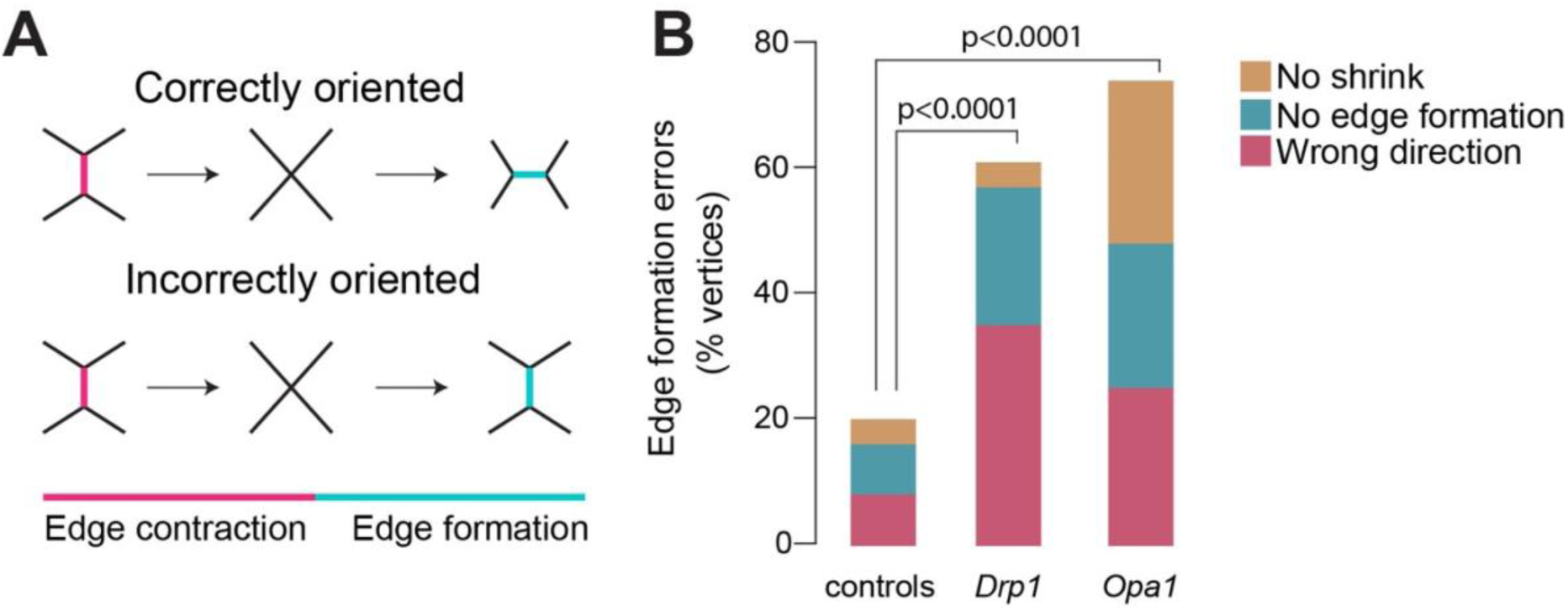
Inhibition of mitochondrial fusion and fission affects cell intercalation during convergent extension. **A)** Schematic of vertical cell interfaces (magenta) contracting (X shape) and resulting in a new edge (cyan) either correctly or incorrectly oriented. **B)** Edge formation errors for controls, *Drp1* knockdowns and *Opa1* knockdowns (n=3 embryos per genotype), ∼150 vertices per embryo. Significance determined by Pearson’s 𝛸^2^ square analysis.

### Disrupting mitochondrial dynamics causes significant shifts in cellular bioenergetics as measured by NADH FLIM

To determine whether we could use NADH FLIM to analyze changes in cellular bioenergetics in live *Drosophila* embryos during CE, we used TPEF microscopy coupled with time-correlated single-photon counting to quantify NADH fluorescence lifetime (𝜏_m_) on a per pixel basis. Mean NADH fluorescence lifetime measurements in wild-type embryos were fairly consistent between replicates and across stages 5–8 of development (mean 𝜏_m_ = 1.4 ns; range 𝜏_m_ = 1.24–1.66 ns), and we observed no significant differences between stages by ANOVA analysis (p>0.7, Figure 4A, Supplemental Table 1). Similarly, we observed no significant differences in the contribution of free NADH percent (A1%) across stages (p>0.2, Figure 4F, Supplemental Table 1). These experiments indicate that there are not significant shifts in the bioenergetics of germband cells over the course of *Drosophila* CE.

**Figure 4.**
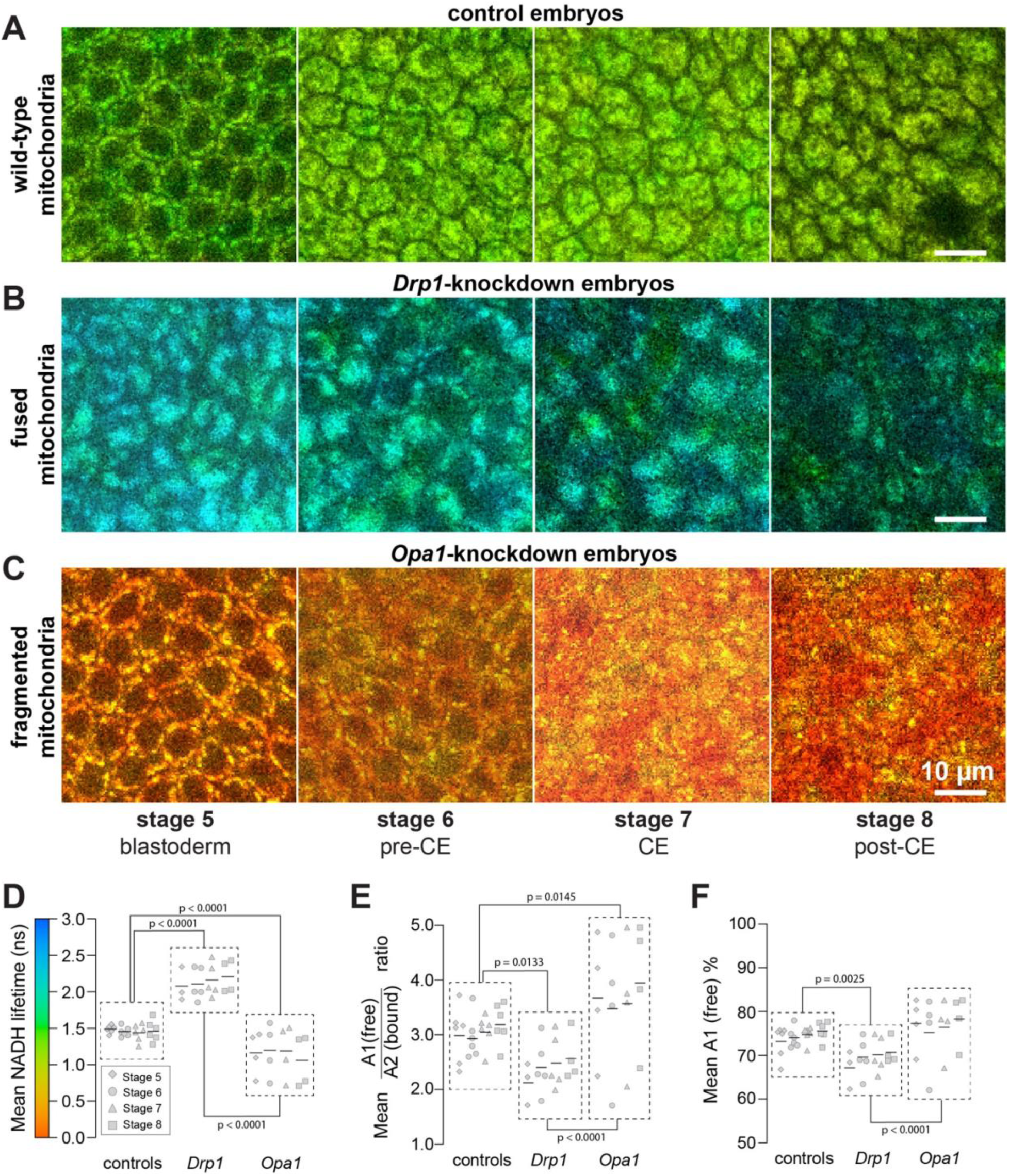
**In vivo fluorescence lifetime imaging (FLIM) quantifies subcellular bioenergetics during *Drosophila* convergent extension**. **A)** Colored maps of NADH FLIM profiles before (stages 5–6), during (stage 7), and after (stage 8) convergent extension in wild-type embryos (n=6). **B)** Fission-impaired embryos (Drp1 knockdown) showed significant differences and dramatic changes in maps (n=3). **C)** Fusion-impaired embryos (Opa1 knockdown) reveal similar FLIM distributions when compared to wild-type, with significant differences in lifetime **(D)** and mean A1/A2 ratio **(E)**. **D)** Summary NADH lifetime comparison of wild-type and disrupted mitochondrial dynamics (n=4). A color scale indicates fluorescence lifetimes, ranging from 0 (red, short lifetimes) to 3 nanoseconds (blue, long lifetimes). **E)** Summary free-to-bound NADH (A1/A2) ratio, comparison of wild-type and disrupted mitochondrial dynamics. **F)** Summary free NADH percent (A1%) comparison of wild-type and disrupted mitochondrial dynamics.

To verify that we could reliably detect shifts in cellular bioenergetics in *Drosophila* embryos, we used FLIM to measure NADH lifetime in *Drp1-*knockdown embryos using weak and strong maternal Gal4 drivers. We predicted that we would see longer NADH lifetimes in these embryos, and that the magnitude of the shifts should correlate with the strength of the Gal4 driver. Using a relatively weak “13-4” maternal Gal4 driver to knock down *Drp1*, we saw a trend towards increased NADH fluorescence lifetimes when considering the data from all stages combined (mean NADH 𝜏_m_ = 1.61 ns vs. 1.4 ns in wild type; range 𝜏_m_ = 1.37–1.93 ns), although these changes were not significantly different from controls at any stage (p>0.2, Supplemental Table 1); the same was true when analyzing A1% (p>0.8, Supplemental Table 1). We did note that mitochondria appeared more clustered during stage 6 compared with controls, and NADH lifetime was significantly lower during stage 8 compared with stage 5 (p = 0.033) (Supplemental Figure 1 and Supplemental Table 1). By contrast, using a relatively strong “67;15” Gal4 driver (same as in Figure 2) to knock down *Drp1*, we observed significant increases in mean NADH lifetime relative to controls for all stages (see Supplemental Figure 1 and Supplemental Table 1 for details), and NADH autofluorescence revealed clear changes in mitochondrial network architecture (Figure 4B,D), similar to those described using conventional immunofluorescence (Figure 2). Like the controls, we did not detect significant differences in NADH lifetime between stages for the strong *Drp1* knockdowns (Supplemental Table 1). Therefore, to simplify our analyses, we pooled the data from all stages for both controls and the strong knockdowns, which yielded a highly significant increase in mean NADH lifetime in *Drp1*-knockdown embryos relative to controls (mean 𝜏_m_ = 2.12 ns vs. 1.4 ns in wild type; p < 0.0001) (Figure 4D). Using the pooled data method, we also saw significant differences for mean A1/A2 ratio (p = 0.0133) (Figure 4E) and mean A1% (p = 0.0025) relative to controls (Figure 4F). These results indicate that FLIM can reliably detect changes in NADH lifetime between fission-inhibited and control *Drosophila* embryos, and that these effects are dose-dependent.

To verify that we could also detect shifts in cellular bioenergetics by inhibiting fusion, we used FLIM to measure NADH lifetime in *Opa1*-knockdown embryos. Knocking down Opa1 using the relatively weak “13-4” maternal Gal4 driver had no significant effects on mitochondrial network morphology or NADH lifetime (data not shown). Knocking down Opa1 using the relatively strong “67;15” maternal Gal4 driver led to hyper-fragmented mitochondrial networks (Figure 4C), similar to what was described in Figure 2. For this genetic background, we did not observe statistically significant differences in NADH lifetime relative to controls for the individual stages, with the exception of stage 8 (p = 0.0350) (Supplemental Table 1), and we also didn’t see significant differences between stages among the *Opa1*-knockdown embryos (Supplemental Table 1). However, when we pooled the data from all stages together, we saw a highly significant decreased in NADH lifetime in *Opa1-*knockdown embryos relative to controls (mean 𝜏_m_ = 1.13 ns vs. 1.4 ns in wild-type; p < 0.0001). We also observed a significant difference relative to controls for mean A1/A2 ratio (p = 0.0145) (Figure 4E), but not for mean A1% (Figure 4F). These results indicate that FLIM can detect changes in NADH lifetime between fusion-inhibited and control *Drosophila* embryos, although the changes may harder to detect compared with inhibiting fission.

## DISCUSSION

In this study, we set out to determine the role of mitochondrial localization and dynamics in a developmental model of epithelial remodeling, namely CE of the *Drosophila* germband epithelium. *Drosophila* CE involves the coordinated intercalation of thousands of epithelial cells to rapidly elongate the germband tissue along the head-to-tail axis. We hypothesized that we might observe significant changes in mitochondrial subcellular localization, network architecture and/or bioenergetics (NADH lifetime) as the germband transitioned from a static state to a dynamically rearranging state. In control embryos, we observed a basal to apical movement of mitochondria during the cellularization stage (stage 5), consistent with previous reports (Chowdhary et al., 2017, 2020). By stage 6, most mitochondria had relocated to the apical region of the cell, where they could potentially influence the cytoskeletal and junctional rearrangements that drive cell intercalation. We originally thought that these cells might require increased ATP to fuel CE, which should be observable by a shift towards fused mitochondrial networks and increased NADH lifetime. Mitochondria remained apical during stages 6 (pre-CE), 7 (CE), and 8 (post-CE), and they did not display obvious changes in network architecture over this period (Figure 2), at least not at the resolution of conventional confocal microscopy. In addition, using FLIM microscopy, we did not observe significant changes in NADH lifetime during stages 5 through 8 in control embryos (Figure 3). These findings suggest that the levels of ATP generation before CE are sufficient to fuel CE without need for significant changes in cellular metabolism.

We were able to use the Gal4/UAS system to drive RNAi to disrupt mitochondrial dynamics in the germband epithelium. In fission-defective *Drp1*-knockdown embryos, mitochondrial networks were noticeably more threadlike, consistent with a hyper-fused state, and FLIM measurements showed a significant shift towards longer NADH lifetime, consistent with more protein-bound NADH and oxidative phosphorylation. By contrast, in fusion-defective *Opa1*-knockdown embryos, mitochondrial networks were more punctate, consistent with a hyper-fragmented state, and FLIM measurements showed a significant shift towards shorter NADH lifetimes, consistent with more cytoplasmic NADH and glycolysis. These results demonstrate that we can reliably detect differences in mitochondrial network architecture and cellular bioenergetics using these experimental approaches. Furthermore, the fact that we can shift mitochondrial dynamics in both directions indicate that wild-type networks are neither hyper-fused nor hyper-fragmented, but rather in an intermediate state during CE.

Notably, we show that disrupting either mitochondrial fission or fusion led to significantly more errors during cell intercalation, demonstrating that mitochondrial dynamics are required for proper CE in *Drosophila*. However, we hypothesize that disrupting fission and fusion interfere with CE for different reasons. In *Drp1-*knockdown embryos, hyper-fused mitochondrial networks are unable to efficiently travel apically along microtubules and remain basal for a significantly longer time than in control embryos (Chowdhary et al., 2017, 2020). Assuming hyper-fused networks can carry out oxidative phosphorylation more efficiently than wild-type networks, we hypothesize that the defects observed in *Drp1*-knockdown embryos are caused by improper subcellular localization of mitochondria and the inability of ATP and/or ROS molecules to diffuse to the apical region of cell. By contrast, in *Opa1-*knockdown embryos, hyper-fragmented networks are present in the apical cell region during stages 6 through 8, and show a similar subcellular localization pattern to control embryos. Assuming hyper-fragmented networks cannot carry out oxidative phosphorylation as efficiently as wild-type networks, we hypothesize the intercalation defects observed in *Opa1*-knockdown embryos are caused by insufficient levels of ATP or ROS molecules, as recently demonstrated in neighboring mesodermal tissues (Madan et al., 2025).

We also wished to determine how the NADH mean lifetime measurements we observed (1.4 ns for control embryos; 2.12 ns for fission-inhibited embryos; and 1.13 ns for fusion-inhibited embryos) in the early embryo compare to those reported in other *Drosophila* studies. We could only find a limited number of other *Drosophila* studies in which multiphoton FLIM was used to measure NADH lifetime in unlabeled samples. Wetzker & Reinhardt (2019) measured mean NADH lifetime in larval salivary glands (1.39 ns), larval fat body (1.68 ns), larval enterocytes (1.06 ns), and adult sperm cells (0.78 ns). They also assessed the percentage of free NADH (A1%) per tissue, reporting 71.2% in larval salivary glands, 61.9% in larval fat body, 72.9% in larval enterocytes, 79.2% in male-stored adult sperm cells, and 78.2% in female-stored adult sperm cells. They concluded that the low mean NADH lifetime and high A1% observed in sperm cells indicated that they were relatively glycolytic compared with somatic cells. Roussel and colleagues (2025) used NADH FLIM to study the role of cellular metabolism in the fly brain. They found that the neuropil region had reduced mean NADH lifetimes (∼1.0-1.3 ns) and higher A1% (∼70%) compared with the brain cortex (mean lifetime = ∼2.5-3.0 ns, A1% = ∼60%). Furthermore, they assessed whether memory formation alters the metabolic signatures of the specific cell types, and they found that mushroom bodies and kenyon cell subtypes α/β (associated with long-term memory) and γ (linked to short-term memory) showed significant differences in their basal A1 percentages (∼73% and ∼70%, respectively). Following paired training tests in adult flies, they observed a shift in A1% for α/β (∼72%) and γ (∼69%) cells in conditioned flies. Notably, our observed A1 percentages (74.2% in controls, 69.5% in fission-inhibited embryos, and 76.7% in fusion-inhibited embryos) were similar to values reported in these studies, indicating that the metabolic characteristics of germband cells undergoing CE are within the range of other somatic tissues in *Drosophila*.

Finally, we wish to address how our results fit in with the many mammalian studies that used NADH FLIM to distinguish cancerous from noncancerous cells. This is possible due to the Warburg effect, in which cancerous cells shift their metabolism to utilize more glycolysis (even in the presence of sufficient oxygen), which leads to more free NADH and shorter NADH lifetimes (Lunt & Vander Heiden, 2011; Yao et al., 2019). Skala et al. used FLIM to characterize epithelial pre-cancer *in vivo*, reporting lifetimes of 2.35 ns in normal tissues and shorter lifetimes in low- and high-grade precancerous animals (2.25 ns and 2.15 ns, respectively). Szulczewski et al. reported an increase in the proportion of free NADH (A1%), leading to short NADH lifetimes for tumor cells in breast tissue, (0.06 – 1.8 ns). Moreover, FLIM analyses have also been reported in clinical studies. Wang et al. demonstrated differences in NADH lifetimes between healthy tissue (1.48 ns) and lung carcinomas (1.22 ns) and suggested the *in vivo* use of FLIM as a non-invasive, highly accurate approach in cancer detection and diagnosis. Consistent trends were also described in tumor-associated macrophages, where low NADH mean lifetimes were observed and associated with glycolytic signatures. Similarly, studies in 3D-engineered *in vitro* cancer models demonstrated a behavior comparable to the Warburg effect in human precancerous epithelial tissue. Furthermore, studies in rat fibrosarcoma W31 cells validated the shifts between free and bound NADH as the source of the observed changes in NADH lifetime, independent of variations in the microenvironment (e.g., pH) or refractive index (Datta et al., 2020; Kolenc & Quinn, 2019). Therefore, we conclude that the NADH FLIM measurements we observed in *Drosophila* embryos also fall within the general range seen for healthy and cancerous mammalian cells, making direct comparisons between *Drosophila* and mammalian studies plausible.

## ACKNOWLEDGEMENTS

We would like to acknowledge generous funding from the NIH NIGMS to A. Paré that directly supported this research (1R01GM147372-01), as well as funding and support from the Arkansas Integrative Metabolic Research Center (AIMRC) (1P20GM139768-01 5743), an NIH Center of Biomedical Research Excellence. Stocks obtained from the Bloomington Drosophila Stock Center (NIH P40OD018537) were used in this study. We would also like to acknowledge FlyBase (http://flybase.org).

---

**Supplemental Figure 1.**
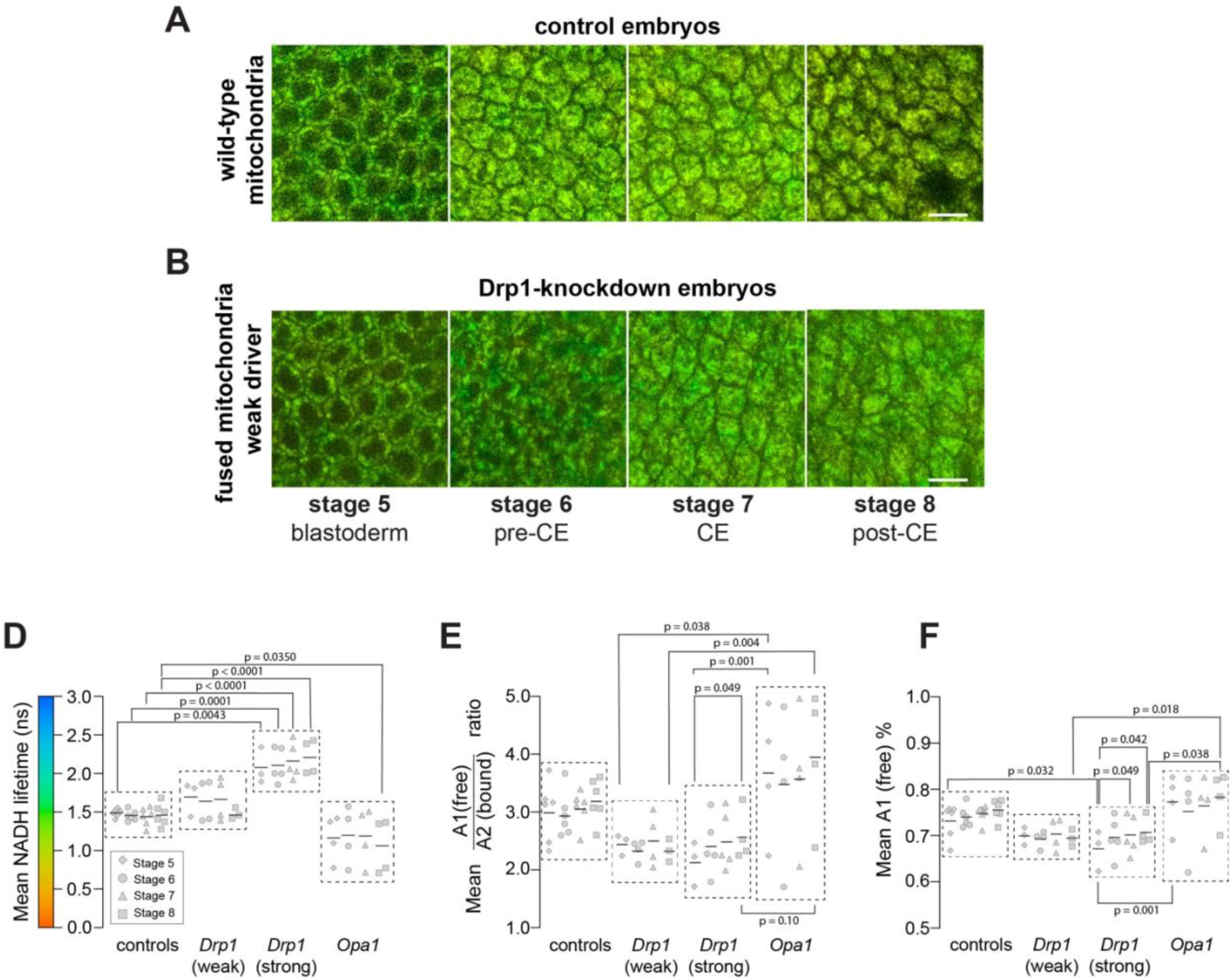
N**A**DH **fluorescence lifetime imaging identifies metabolic shifts during *Drosophila* convergent extension**. Spatial distribution maps of NADH lifetime before (stages 5–6), during (stage 7), and after (stage 8) convergent extension in **(A)** control embryos and **(B)** fission-impaired embryos (Drp1 knockdown) with a weak driver showed subtle changes in maps (n=3). **D)** Summary NADH lifetime comparison of CE stages between controls and embryos with disrupted mitochondrial dynamics. A color scale indicates fluorescence lifetimes, ranging from 0 ns (red, short lifetimes) to 3 ns (blue, long lifetimes). **E)** Summary mean free-to-bound NADH across genotypes and CE stages. **F)** Summary mean free NADH percentage across genotypes and CE stages.

**Table 1.** ANOVA and Tukey’s multiple comparisons test summary. Mean NADH lifetime, free-to-bound NADH (A1/A2) ratio, and free NADH percent (A1%) significance within stages and among genotypes. Bold values indicate significance. ⍺ = 0.05.

|  | Mean NADH Lifetime | Mean free-to-bound (A1/A2) ratio | Mean free NADH percent (A1 %) |
| --- | --- | --- | --- |
|  | Adjusted P Value | Adjusted P Value | Adjusted P Value |
| **Control** |  |  |  |
| Stage 5 vs. Stage 6 | 0.9041 | 0.988 | 0.922 |
| Stage 5 vs. Stage 7 | 0.7792 | 0.978 | 0.62 |
| Stage 5 vs. Stage 8 | 0.9656 | 0.627 | 0.281 |
| Stage 6 vs. Stage 7 | 0.9931 | 0.886 | 0.935 |
| Stage 6 vs. Stage 8 | 0.9966 | 0.43 | 0.64 |
| Stage 7 vs. Stage 8 | 0.9631 | 0.852 | 0.932 |
| **Drp1 KD (weak)** |  |  |  |
| Stage 5 vs. Stage 6 | 0.2675 | 0.994 | >.999 |
| Stage 5 vs. Stage 7 | 0.4427 | 0.953 | 0.964 |
| Stage 5 vs. Stage 8 | **0.033** | 0.959 | 0.993 |
| Stage 6 vs. Stage 7 | 0.9814 | 0.823 | 0.916 |
| Stage 6 vs. Stage 8 | 0.6846 | 0.993 | 0.998 |
| Stage 7 vs. Stage 8 | 0.4765 | 0.727 | 0.873 |
| **Drp1 KD (strong)** |  |  |  |
| Stage 5 vs. Stage 6 | 0.4858 | 0.123 | 0.097 |
| Stage 5 vs. Stage 7 | 0.1373 | 0.056 | **0.049** |
| Stage 5 vs. Stage 8 | 0.0691 | **0.049** | **0.042** |
| Stage 6 vs. Stage 7 | 0.7844 | 0.971 | 0.983 |
| Stage 6 vs. Stage 8 | 0.5745 | 0.949 | 0.962 |
| Stage 7 vs. Stage 8 | 0.9735 | >.999 | 0.999 |
| **Opa1 KD (strong)** |  |  |  |
| Stage 5 vs. Stage 6 | 0.9635 | 0.763 | 0.569 |
| Stage 5 vs. Stage 7 | 0.985 | 0.953 | 0.947 |
| Stage 5 vs. Stage 8 | 0.3038 | 0.529 | 0.926 |
| Stage 6 vs. Stage 7 | 0.9992 | 0.969 | 0.875 |
| Stage 6 vs. Stage 8 | 0.1286 | 0.106 | 0.238 |
| Stage 7 vs. Stage 8 | 0.165 | 0.249 | 0.653 |
| **Stage 5** |  |  |  |
| Control vs. Drp1 KD (weak) | 0.251 | 0.5 | 0.555 |
| Control vs. Drp1 KD (strong) | **0.0043** | 0.054 | **0.032** |
| Control vs. Opa1 KD | 0.1412 | 0.361 | 0.421 |
| Drp1 KD (weak) vs. Drp1 KD (strong) | 0.4994 | 0.741 | 0.574 |
| Drp1 KD (weak) vs. Opa1 KD | **0.003** | **0.038** | 0.06 |
| Drp1 KD (strong) vs. Opa1 KD | **<0.0001** | **0.001** | **0.001** |
| **Stage 6** |  |  |  |
| Control vs. Drp1 KD (weak) | 0.5961 | 0.471 | 0.309 |
| Control vs. Drp1 KD (strong) | **0.0001** | 0.534 | 0.321 |
| Control vs. Opa1 KD | 0.321 | 0.564 | 0.967 |
| Drp1 KD (weak) vs. Drp1 KD (strong) | **0.0195** | 0.998 | >.999 |
| Drp1 KD (weak) vs. Opa1 KD | **0.0423** | 0.068 | 0.194 |
| Drp1 KD (strong) vs. Opa1 KD | **<0.0001** | 0.075 | 0.201 |
| **Stage 7** |  |  |  |
| Control vs. Drp1 KD (weak) | 0.4343 | 0.552 | 0.36 |
| Control vs. Drp1 KD (strong) | **<0.0001** | 0.472 | 0.275 |
| Control vs. Opa1 KD | 0.3417 | 0.606 | 0.924 |
| Drp1 KD (weak) vs. Drp1 KD (strong) | **0.0114** | >.999 | >.999 |
| Drp1 KD (weak) vs. Opa1 KD | **0.0252** | 0.102 | 0.174 |
| Drp1 KD (strong) vs. Opa1 KD | **<0.0001** | 0.071 | 0.125 |
| **Stage 8** |  |  |  |
| Control vs. Drp1 KD (weak) | 0.9271 | 0.154 | 0.1 |
| Control vs. Drp1 KD (strong) | **<0.0001** | 0.339 | 0.206 |
| Control vs. Opa1 KD | **0.035** | 0.271 | 0.736 |
| Drp1 KD (weak) vs. Drp1 KD (strong) | **0.0013** | 0.95 | 0.963 |
| Drp1 KD (weak) vs. Opa1 KD | **0.0172** | **0.004** | **0.018** |
| Drp1 KD (strong) vs. Opa1 KD | **<0.0001** | **0.01** | **0.038** |

